# Functional insights from KpfR, a new transcriptional regulator of fimbrial expression that is crucial for *Klebsiella pneumoniae* pathogenicity

**DOI:** 10.1101/2020.08.31.276717

**Authors:** Ana E. I. Gomes, Thaisy Pacheco, Cristiane S. Santos, José A. Pereira, Marcelo L. Ribeiro, Michelle Darrieux, Lúcio F. C. Ferraz

**Affiliations:** Laboratory of Molecular Biology of Microorganisms, Sao Francisco University, Braganca Paulista, Sao Paulo, Brazil; Cellular and Molecular Biology of Tumors, Sao Francisco University, Braganca Paulista, Sao Paulo, Brazil; Laboratory of Immunopharmacology and Molecular Biology, Sao Francisco University, Braganca Paulista, Sao Paulo, Brazil

**Author notes:** Address correspondence to Lucio F. C. Ferraz. These authors contributed equally to this work (Ana E. I. Gomes, Thaisy Pacheco).

**Keywords:** *Klebsiella pneumoniae*, transcriptional regulation, fimbriae, adherence, biofilm, coculture, urinary tract infection, host-microbe interaction

## Abstract

Although originally known as an opportunistic pathogen, *Klebsiella pneumoniae* has been considered a worldwide health threat nowadays due to the emergence of hypervirulent and antibiotic-resistant strains capable of causing severe infections not only on immunocompromised patients but also on healthy individuals. Fimbriae is an essential virulence factor for *K. pneumoniae*, especially in urinary tract infections, because it allows the pathogen to adhere and invade urothelial cells and to form biofilms on biotic and abiotic surfaces. The importance of fimbriae for *K. pneumoniae* pathogenicity is highlighted by the large number of fimbrial gene clusters on the bacterium genome, which requires a coordinated and finely adjusted system to control the synthesis of these structures. In this work, we describe KpfR as a new transcriptional repressor of fimbrial expression in *K. pneumoniae* and discuss its role in the bacterium pathogenicity. *K. pneumoniae* lacking the *kpfR* gene exhibited a hyperfimbriated phenotype with enhanced biofilm formation and greater adhesion to and replication within epithelial host cells. However, the mutant strain was attenuated for colonization of the bladder in a murine model of urinary tract infection. These results indicate that KpfR is an important transcriptional repressor that, by negatively controlling the expression of fimbriae, prevents *K. pneumoniae* from having a hyperfimbriated phenotype and from being recognized and eliminated by the host immune system.

**IMPORTANCE:** Fimbriae are crucial virulence factor for many pathogens because they allow the pathogen to adhere and invade host cells and to form biofilm on biotic and abiotic surfaces. However, the synthesis of fimbriae requires a precise and coordinated control to guarantees the production of these structures only when necessary, thus avoiding unnecessary energy expenditure and bacterial clearance by the host immune cells. Herein, we describe for the first time the role of the transcriptional repressor of fimbrial expression KpfR on the pathogenicity of *K. pneumoniae*, a Gram-negative pathogen that has gained worldwide attention, notably for being the causative agent of severe and metastatic infections even on healthy individuals. By deleting the *kpfR* gene, we show that the mutant strain loses the control of fimbriae production, resulting in a hyperfimbriated phenotype that impairs *K. pneumoniae* ability to colonize a murine model of urinary tract infection.

## INTRODUCTION

*Klebsiella pneumoniae* is a Gram-negative pathogen responsible for a wide range of healthcare-acquired infections in genitourinary, respiratory and gastrointestinal tracts, mostly in immunocompromised patients (1, 2). Polysaccharide capsule, lipopolysaccharide, siderophore-mediated iron acquisition systems and fimbrial adhesins are among the virulence determinants of *K. pneumoniae* (2). Currently, this bacterium has become a worldwide public health threat due to the emergence of hypervirulent and antibiotic-resistant strains of *K. pneumoniae* causing various types of invasive infections (3-5). In addition, the increasing number of severe community-acquired infections in healthy individuals has emphasized the importance of studying the virulence mechanisms that determine the pathogenicity of *K. pneumoniae*.

Although historically known for being responsible for pneumonia, *K. pneumoniae* has been considered an important causative agent of urinary tract infections (UTIs) (2). The successful colonization of the urinary epithelium by *K. pneumoniae* relies on the expression of fimbriae. These structures mediate adherence to urothelial cells and are also essential for biofilm formation by promoting adhesion of the bacterium to abiotic surfaces, such as urinary catheters. The establishment of a biofilm structure renders the bacterium more resistant to the host defense system and increases colonization of the host urinary tract (6-12). Fimbriae are also required for uropathogens to adhere to specific receptors on urothelial cells and invade the urinary epithelium. Once in the intracellular environment, the uropathogen replicates to form biofilm-like intracellular bacterial communities (IBCs) (13). Later maturation of IBCs leads to epithelium exfoliation with consequent spread of the bacteria to other sites, initiating a new cycle (6, 13). Thus, this intracellular niche allows uropathogens to hide from the host immune system and represents a quiescent intracellular reservoir of bacteria that leads to the recurrent episodes of UTI (6, 13).

Several types of fimbriae have been identified in *K. pneumoniae*, with types 1 and 3 fimbriae being the main and best characterized adhesive structures (2). Type 1 fimbriae are encoded by the *fim* gene cluster, composed of seven structural genes (*fimAICDFGH*). These fimbriae are made up of repeating subunits of the major fimbrial subunit protein FimA, with the adhesin molecule FimH located at the distal end of the structure. FimH has a binding affinity for mannose residues present on bladder cells surface. Thus, type 1 fimbriae promote the adhesion and invasion of the pathogen in host cells and favor a successful colonization (7, 9). Type 3 fimbriae are encoded by the *mrkABCD* gene cluster. MrkA subunits make up the major structure of the type 3 fimbriae, with the MrkD adhesin located at the tip. Although the MrkD-specific cell surface receptor has not yet been identified, type 3 fimbriae promote adhesion to tracheal, lung and kidney epithelial cells (14), and are fundamental in the process of biofilm formation on biotic and abiotic surfaces (15). Both type 1 and type 3 fimbriae promote biofilm formation on urinary catheters and, therefore, play a significant role in colonization and persistence in the bladder in catheter-associated UTI (11, 12).

*K. pneumoniae* controls the expression of fimbrial genes through transcriptional regulators, utilizing various environmental stimuli and according to specific anatomic sites. Phase variation mediated by invertible DNA elements (9, 16), intracellular levels of cyclic di-GMP (17) and DNA binding regulators (7, 18, 19) are among the known mechanisms that regulate expression of fimbriae in *K. pneumoniae* (20). For instance, the expression of type 3 fimbriae has been attributed to extracellular iron levels and ferric uptake regulator (Fur) (21). Fur is a transcriptional regulator that modulates gene expression by complexing with ferrous iron and binding to regulatory sequences, named boxes Fur, located in the promoter region of target genes (22). On the other hand, the expression of *fim* gene cluster is regulated by phase variation mediated by an invertible promoter element, named *fimS* element, whose orientation is switched by FimE- and FimB-recombinases (9). Besides this mechanism, the expression of *K. pneumoniae fim* gene cluster is also regulated by the transcriptional regulator FimK encoded by *fimK*, a gene found only in *fim* gene cluster of *K. pneumoniae* and absent in *Escherichia coli* (7, 19).

In addition to *fim* and *mrK*, the genome of *Klebsiella pneumoniae* presents at least seven other fimbrial gene clusters still poorly characterized (16, 23). One of the clusters not yet characterized is *kpf* gene cluster, first described by Wu and colleagues on *K. pneumoniae* NTUH-K2044 (16). The synthesis of this broad repertoire of fimbrial structures requires a precise and coordinated control that involves specific regulatory proteins. This control guarantees the production of these structures only when necessary, avoiding uncontrolled expression of fimbriae and unnecessary energy expenditure by bacteria. Besides resulting in unnecessary consumption of energy, the uncontrolled expression of the fimbriae may result in bacterial clearance by the immune cells (24, 25).

In the present study, we describe the transcriptional regulator of *kpf* gene cluster, a poorly characterized cluster of fimbrial genes encoding type 1-like fimbriae. We also demonstrate that KpfR regulator plays an important role in the pathogenicity of *K. pneumoniae* in murine urinary tract.

## RESULTS

### *kpfR* is a Fur-regulated gene that encodes the transcriptional regulator of *kpf* gene cluster

The *kpf* gene cluster comprises 4 genes designated as follow: *kpfA*, encoding the major pilin subunit, *kpfB*, encoding a chaperone, *kpfC*, encoding an usher, and *kpfD*, encoding an adhesin. The first gene adjacent to *kpf* cluster at 5’ extremity encodes a putative helix-turn-helix transcriptional regulator. To determine whether this adjacent gene belongs to *kpf* cluster, reverse-transcription-PCRs were performed using cDNA and primer pairs spanning the entire cluster and confirmed that *kpf* cluster and the adjacent gene are co-transcribed as a single polycistronic transcript (Fig. 1A). Since the adjacent gene encodes a putative helix-turn-helix transcriptional regulator, and following Wu et al. nomenclature, we designated it *kpfR* gene (16).

**Figure 1.**
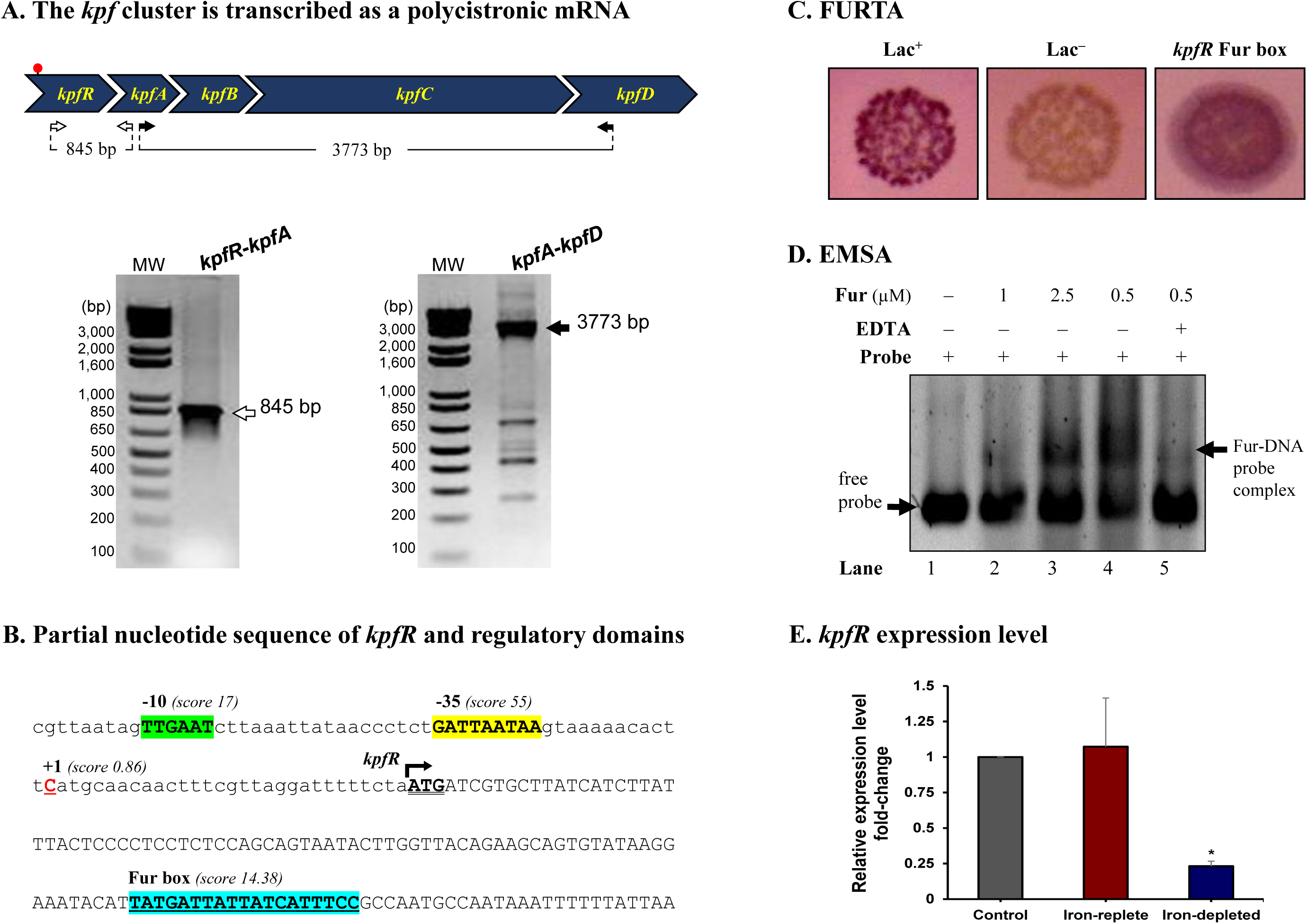
The *kpf* gene cluster encoding type 1-like fimbriae is regulated by Fur. (A) *kpf* cluster comprises *kpfR, kpfA, kpfB, kpfC* and *kpfD* genes and is transcribed as a polycistronic mRNA. cDNA synthesized from total RNA of *K. pneumoniae* was used in PCR reactions using the primer pairs represented in the scheme. The resulting amplicons were analyzed by agarose gel electrophoresis and confirmed the predicted sizes of 847 and 3773 base pairs (bp). The red spot on the scheme indicates the putative Fur box motif identified on *kpf* cluster. (B) Partial sequence of *kpfR* showing the initial codon (ATG, double underlined) and the Fur-binding sequence located inside the coding region of *kpfR* (highlighted in blue). Also indicated on the promoter region of the cluster are the -10 (green) and -35 (yellow) domains of the housekeeping Sigma factor 70 and the predicted transcription initiation site at position +1 (cytosine in red). (C to D) Fur Titration Assay (FURTA) and DNA Electrophoretic Mobility Shift Assay (EMSA) confirm that Fur protein recognizes and binds to the Fur box on *kpfR* of *K. pneumoniae*. On FURTA, the Fur box was validated, as indicated by the red colonies similar to the FURTA-positive control (Lac^+^). On EMSA, Fur-DNA probe are complexed as increased concentrations of *K. pneumoniae* purified His-Fur protein were incubated with the probes, thus confirming the direct interaction of *K. pneumoniae* Fur protein on the putative Fur box found on *kpfR*. Fur interaction depends on divalent cations, since the addition of EDTA chelator abolished the mobility shift of the probe. (E) RT-qPCR analyses showed that Fur represses the expression of the *kpfR* gene on *K. pneumoniae* cells cultured under iron-depleted condition when compared to the control condition (bacteria cultured in LB medium only). Since *kpfR* belongs to the *kpf* gene cluster, Fur modulates the expression of the entire cluster according to the availability of iron in the culture medium. *, *p*-value ≤ 0.05.

To better understand the expression regulation of *kpf*, bioinformatic analyses were employed to identify regulatory domains in the promoter region of the cluster. Thus, a putative Fur box motif was identified inside the coding region of *kpfR* (Fig. 1B). This putative Fur box was validated by Fur Titration Assay (FURTA) and DNA Electrophoretic Mobility Shift Assay (EMSA). Fig. 1C shows the FURTA-positive control (Lac^+^, red colonies), the FURTA-negative control (Lac^−^, colorless colonies) and the validation of the Fur box on *kpfR*, indicated by the red colonies. This result confirms that Fur regulator from *E. coli* H1717 was able to bind *in vivo* to the cloned putative Fur box from *kpfR* of *K. pneumoniae*, rendering the colonies with red color.

Next, EMSA was performed to verify the direct interaction of *K. pneumoniae* Fur protein on the putative Fur box found on *kpfR*. As shown on Fig. 1D, a mobility shift of the DNA probes containing the putative Fur box was observed when increasing concentrations of *K. pneumoniae* purified His-Fur protein was incubated with these probes. Besides, the addition of the divalent cations chelator EDTA abolished the mobility shift of the probe, indicating that divalent cations are required for Fur interaction.

In summary, our results confirm that Fur protein recognizes and binds to the Fur box on *kpfR* of *K. pneumoniae*, indicating that the expression of *kpfR* seems to be modulated by Fur transcriptional regulator in an iron-dependent manner. To further investigate how Fur modulates the expression of *kpfR*, Reverse Transcription Quantitative real-time Polymerase Chain Reaction (RT-qPCR) analyses were performed on *K. pneumoniae* cells kept under iron-replete and iron-limited conditions. As displayed on Fig. 1E, the iron-replete condition presents no changes in the expression pattern of *kpfR* gene when compared to the control condition. On the other hand, *kpfR* is repressed under condition of iron scarcity. These results show that Fur modulates the expression of *kpfR* gene according to the availability of iron in the culture medium. Thus, *kpfR* belongs to the Fur regulon of *Klebsiella pneumoniae*.

### *kpfR* disruption modified the bacterial morphology and enabled a rudimentary motility

To evaluate the role of KpfR as regulator of morphology and virulence of *Klebsiella pneumoniae*, a mutant (*kpfR*::*kan*^R^) was generated (as described in Methods). The mutant strain exhibits prolonged growth at lag phase, leading to delayed entry into logarithmic (log) phase of growth (see Fig. S1 in the supplemental material). However, once reaching the log phase, the mutant strain displayed a growth rate similar to that observed in the wild-type strain. Slight differences in growth patterns were observed among wild-type and complemented strains.

Macroscopic morphologies of wild-type UKP8 and mutant *kpfR*::*kan*^R^ strains were investigated on blood agar plates. As shown on Fig. 2A, the wild-type strain exhibited circular shape, high elevation, entire margin and smooth bright surface. On the other hand, the mutant *kpfR*::*kan*^R^ strain presented irregular shape, flat elevation, lobed margin and rough matte surface. In addition, *kpfR*::*kan*^R^ colonies exhibited a slight dispersion on the plates. The morphological features observed on the mutant strain can be attributed to the lack of KpfR, because the pattern observed on the wild-type strain was reestablished on the complemented mutant strain (*kpfR*::*kan*^R^_comp_).

**Figure 2.**
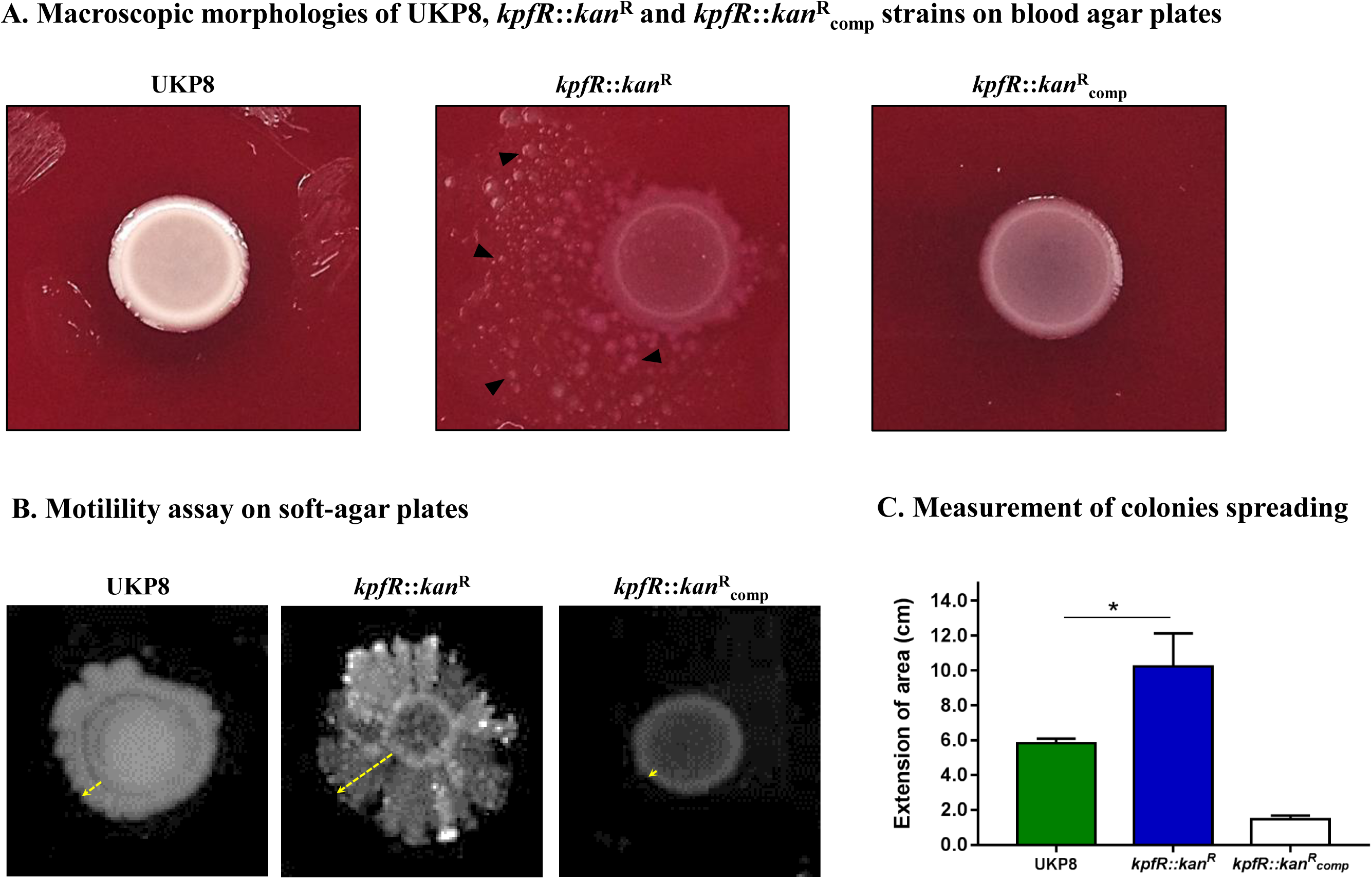
Bacterial colonies of *K. pneumoniae* cells depleted of *kpfR* show altered macroscopic morphologies and rudimentary motility. (A) Colony morphology of wild-type (UKP8), mutant (*kpfR*::*kan*^R^), and complemented (*kpfR*::*kan*^R^_comp_) strains were investigated on blood agar plates. The altered pattern on the mutant strain was reverted in the complemented strain, which exhibited a pattern similar to the wild-type. (B and C) In soft agar plates, *kpfR*::*kan*^R^ cells exhibited a pronounced asymmetric dispersion on the surface of the plates, resembling a sort of rudimentary motility. This greater dispersion of the mutant colonies than the wild-type is statistically significant. *, *p*-value ≤ 0.05.

The spreading behavior of *kpfR*::*kan*^R^ colonies on blood agar plates prompted us to perform motility assays on soft agar plates. As displayed on Fig. 2B, the wild-type UKP8 strains presented a slight and linear radial migration on soft agar plates. On the other hand, *kpfR*::*kan*^R^ cells exhibited a more prominent and asymmetric dispersion on the surface of the plates. The complemented mutant strain had a reduced migration extension when compared to both wild-type and mutant strains. The extent of the area in which the wild-type, mutant and complemented strains migrated on the plates was measured, and confirmed that the mutant strain had a greater dispersion when compared to UKP8 and *kpfR*::*kan*^R^_comp_ strains (Fig. 2C). These results indicate that *kpfR* disruption rendered to the mutant bacteria a sort of rudimentary motility.

### Lack of KpfR triggers the production of bacterial surface appendages, enhances yeast agglutination and biofilm formation but reduces capsule production

Since loss of *kpfR* rendered *K. pneumoniae* a prominent dispersion on agar plates, we evaluated the cell surface morphology of the wild-type, mutant and complemented strains by Transmission Electron Microscopy (TEM). While UKP8 strain appears with no surface appendages, the cell surface of the *kpfR*::*kan*^R^ cells were covered by numerous irregular appendages similar to fimbriae or pili that are long and flexible with a varied size larger than 1 μm (Fig. 3A). These numerous appendages were no longer seen on the surface of *kpfR*::*kan*^R^_comp_ strain.

**Figure 3.**
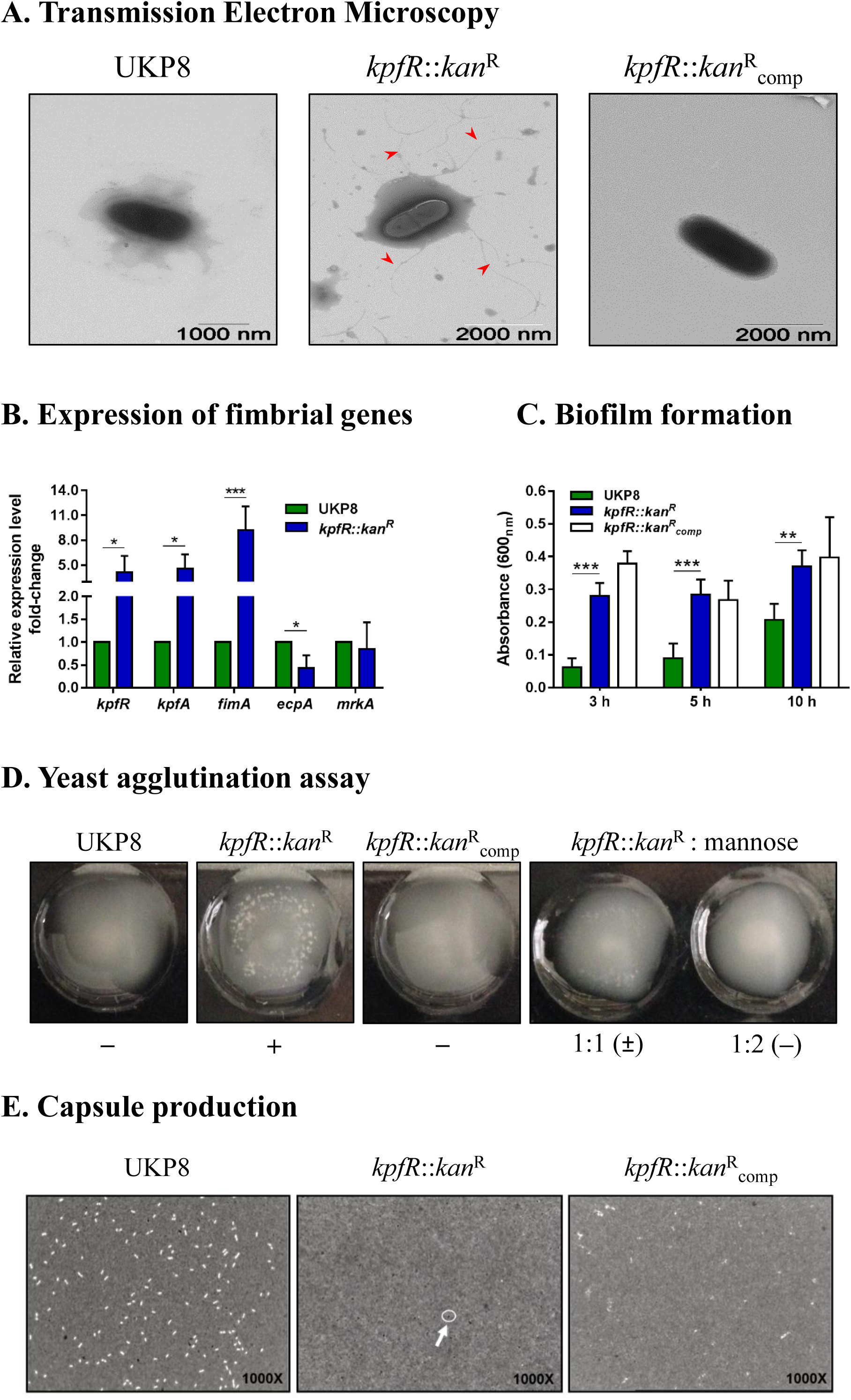
Lack of KpfR triggers the expression of type 1-like fimbriae, increases biofilm formation and yeast cells agglutination, and reduces production of capsule polysaccharide. (A) TEM analyses revealed *kpfR*::*kan*^R^ mutant cells covered with numerous fimbriae-like appendages that were absent in wild-type and complemented *kpfR*::*kan*^R^_comp_ cells. (B) This hyperfimbriated phenotype is corroborated by the fimbrial genes expression analyses. Although *ecpA*, from *ecp* gene cluster, is downregulated, the *kpfR*::*kan*^R^ mutant cells present up-regulation of *fimA* (*fim* gene cluster of type 1 fimbriae) and *kpfR* and *kpfA* genes (*kpf* gene cluster of type 1-like fimbriae). (C and D) Because of the hyperfimbriated phenotype, the mutant strain forms more biofilm than the wild-type and agglutinates yeast cells. This agglutination is mediated by type 1-like fimbriae because in the presence of mannose the mutant strain loses the ability to agglutinate yeast cells. The images are representative of independent experiments. E) Capsule polysaccharide production is reduced in the *kpfR*::*kan*^R^ mutant strain. Wild-type (UKP8), mutant (*kpfR*::*kan*^R^), and complemented (*kpfR*::*kan*^R^_comp_) strains were cultured under the same growth conditions, stained with India ink, and analyzed by optical microscopy. The presence of capsule production is indicated by a negative staining area around the bacteria. Unlike what was observed in the wild-type and complemented strains, no visible capsule is observed on *kpfR*::*kan*^R^, suggesting a reduction in capsule biosynthesis on the hyperfimbriated mutant strain. The images are representative of independent experiments. *, *p*-value ≤ 0.05; **, *p*-value ≤ 0.01; ***, *p*-value ≤ 0.005.

The presence of numerous appendages on *kpfR*::*kan*^R^ cells led us to investigate the expression of fimbrial genes on UKP8 and *kpfR*::*kan*^R^ cells grown on blood agar plate. Disruption of *kpfR* resulted in up-regulation of *fimA*, from type 1 fimbrial gene cluster *fim*, and *kpfR* and *kpfA* genes, from *kpf* gene cluster (Fig. 3B). *kpfR* and *kpfA* presented a similar expression pattern, confirming the polycistronic transcription of these genes, as previously shown. On the other hand, lack of KpfR regulator resulted in down-regulation of *ecpA* gene, from the *ecp* cluster, whereas *mrkA* gene, from type 3 fimbrial gene cluster *mrk*, presented unchanged expression pattern between wild-type and mutant strains.

To investigate the phenotypic effects of increased type 1-like fimbriae production due to KpfR absence, biofilm formation and yeast agglutination assays were performed with the UKP8, *kpfR*::*kan*^R^ and *kpfR*::*kan*^R^_comp_ strains. Biofilm formation by *kpfR*::*kan*^R^ cells was significantly superior than UKP8 after 3, 5 and 10 h of incubation (Fig. 3C). The complemented *kpfR*::*kan*^R^_comp_ strain exhibited a phenotype similar to *kpfR*::*kan*^R^. Yeast agglutination assays allow specific detection of type 1 fimbriae because type 1 fimbrial adhesins have great affinity for mannose-containing receptors on the yeast cell-surface. UKP8 showed no yeast agglutination, whereas the *kpfR*::*kan*^R^ promptly agglutinated the yeast cells (Fig. 3D). The lack of agglutination observed on UKP8 was restored on the *kpfR*::*kan*^R^_comp_ strain. To determine if the agglutination pattern exhibited by the mutant strain was due to type 1 fimbriae expression, the assay was also performed in the presence of mannose prior to the addition of the yeast cells. In the presence of mannose, the mutant strain loses the ability to agglutinate yeast cells. These results reveal that the observed agglutination of yeast cells was indeed mediated by type 1 fimbriae.

As the lack of KpfR regulator resulted in increased expression of type 1-like fimbriae, we investigated the production of capsular polysaccharide, another important virulence factor of the bacterial cell surface. Negative staining with India ink revealed that *kpfR*::*kan*^R^ mutant strain presents no visible capsule as compared to the wild-type and complemented strains under the optical microscope (Fig. 3E), suggesting a reduction in capsule biosynthesis on the hyperfimbriated mutant strain.

### *kpfR*::*kan*^R^ colonizes more efficiently bladder epithelial cells *in vitro* but loses resistance *in vivo* and is unable to form IBC in mouse urinary bladder

Considering fimbriae important mediators of bacterial adhesion and internalization into host cells, we investigated the role of *kpfR* gene in host-pathogen interactions by assessing the adhesion, invasion and intracellular replication of UKP8 and *kpfR*::*kan*^R^ strains on human bladder epithelial cell line T24. The data presented on Fig. 4A indicate that the *kpfR*::*kan*^R^ strains showed greater adhesion to T24 bladder cells than the wild-type UKP8 strain. Moreover, only the mutant strain was able to invade and replicate within T24 cells (Fig. 4A). No significant changes in T24 cells viability were observed at a multiplicity of infection (MOI) of 200 bacteria:1 T24 cell and incubation of coculture for 24 h (data not shown). According to the results, the lack of KpfR repressor improved the ability of the mutant strain to adhere, invade and even replicate intracellularly in T24 bladder cells. We next evaluated the expression of *kpfR* and *kpfA* (cluster *kpf*) and *fimA* (cluster *fim*) in UKP8 and *kpfR*::*kan*^R^ strains during adhesion to the T24 cells. As shown on Fig. 4B, all three genes had the expression significantly induced in *kpfR*::*kan*^R^ compared to UKP8.

**Figure 4.**
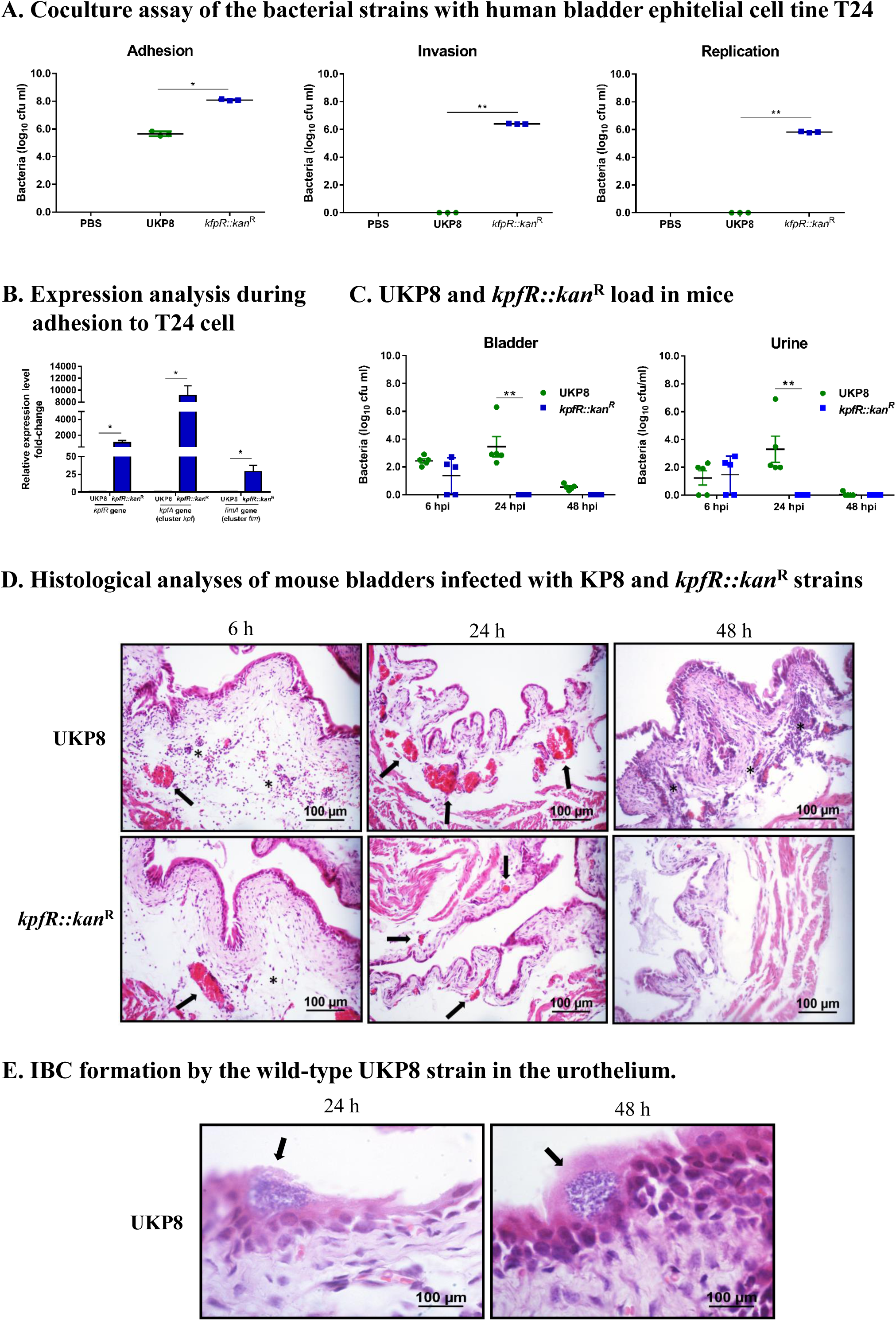
*K. pneumoniae* mutant for *kpfR* adheres more efficiently and replicates within bladder epithelial cells, but loses resistance in the mouse model of urinary infection. For the coculture assays, human bladder epithelial cell line T24 were inoculated with the wild-type and *kpfR*::*kan*^R^ strains at an MOI of 200 to assess the adhesion, invasion, and intracellular replication of the strains on the host human bladder cell. The incubation periods are described in Materials and Methods. (A) The mutant strain has greater adhesion to T24 bladder cells than the wild-type UKP8 strain and is the only strain able to invade and replicate within T24 cells. (B) During adhesion to T24 bladder cells, the *kpfR*::*kan*^R^ mutant strain have increased expression of *kpfR* and *kpfA* (cluster *kpf*) and *fimA* (cluster *fim*) than the wild-type UKP8 strain, which may explain the improved ability of the mutant strain to adhere the T24 cells. *, *p*-value ≤ 0.05; **, *p*-value ≤ 0.01. For the mouse model of urinary infection, animals were inoculated by transurethral catheterization with 5 x 10^8^ CFU/ml of UKP8 and *kpfR*::*kan*^R^ strains. At the indicated times, the animals were euthanized, and urine and bladders were aseptically collected and processed for CFU enumeration and H&E staining. (C) Bacterial CFUs were counted on urine and bladder tissue after 6, 24, and 48 hpi with UKP8 and *kpfR*::*kan*^R^ strains. In mice urine, the wild-type strain is recovered at 6 and 24 hpi, while the mutant *kpfR*::*kan*^R^ strain is recovered only at 6 hpi. In the bladder tissue, the wild-type is present at 6, 24, and 48 hpi, whereas the mutant strain is also recovered only after 6 hpi. Mice are represented by symbols. Data were statistically analyzed by ANOVA test; **, *p*-value ≤ 0.01. (D) Histological analyses of bladders infected with the wild-type and *kpfR*::*kan*^R^ mutant strains show both strains triggering an inflammatory infiltrate consisting of neutrophils (asterisks) and a hyperemia (arrows) at 6 hpi. At 24 hpi, the hyperemia is more pronounced with the wild-type strain, and inflammatory cells migration is observed at 48 hpi only with UKP8, suggesting that at this period of infection the mutant strain has already been completely eliminated. Images are from an individual representative experiment. (E) Only the wild-type UKP8 is able to form biofilm-like intracellular bacterial communities (IBCs) in the urothelium of mice. Images from histological analyses of bladders infected with the wild-type at 24 and 48 hpi reveal IBCs (arrows) within superficial urothelial cells. Images are from an individual representative experiment.

To further evaluate the *in vivo* role of KpfR repressor in *Klebsiella pneumoniae* virulence, a mouse urinary tract infection model was used to compare the ability of UKP8 and *kpfR*::*kan*^R^ strains to colonize the bladder of the animals (Fig. 4C). At 6 hpi, there were no differences in UKP8 and *kpfR*::*kan*^R^ load in mice urine and bladder. At 24 hpi, UKP8 was recovered from bladder tissue and urine, while *kpfR*::*kan*^R^ was not present on these samples. At 48 hpi, UKP8 and *kpfR*::*kan*^R^ were no longer found in urine, whereas only UKP8 was still present in mice bladder tissue.

Aliquots of urine were analyzed under an optical microscope (data not shown) and revealed red blood cells, leukocytes and scaly epithelium in urine of the animals infected with both UKP8 and *kpfR*::*kan*^R^ strains at 6 hpi. At 24 hpi, a larger population of leukocytes and bacteria were observed only in the urine samples of mice infected with the wild-type strain. Urine of mice infected with *kpfR*::*kan*^R^ during 24 h presented neither erythrocytes nor leukocytes.

The colonization experiments suggested that the lack of KpfR repressor confers a disadvantage on *kpfR*::*kan*^R^ strain. To characterize these effects further, histological analyses were performed to obtain images of mouse bladders infected with the wild-type and *kpfR*::*kan*^R^ mutant strains. As shown on Fig. 4D, an inflammatory infiltrate consisting mainly of neutrophils and hyperemia of the bladder blood vessels was observed 6 hpi with both strains. At 24 hpi, the hyperemia was more prominent in the bladder tissue infected with UKP8 than *kpfR*::*kan*^R^, and an abundant migration of inflammatory cells was noticed only with UKP8 at 48 hpi (Fig. 4D). In addition, only wild-type strain was able to form intracellular bacterial communities (IBCs) in the urothelium at 24 and 48 hpi (Fig. 4E). Taken together, the data show successful bladder colonization and formation of IBC in urothelial cells by the wild-type UKP8 strain, while the *kpfR*::*kan*^R^ mutant strain loses resistance and ability to survive in the host.

## DISCUSSION

Fimbriae are important mediators of urinary tract infections caused by *K. pneumoniae* (7, 11, 26, 27). Considering the broad number of fimbrial gene clusters on the *K. pneumoniae* genome, the expression of this range of fimbrial structures demands a coordinated and finely adjusted control. In the current study, we describe the role of KpfR as a transcriptional regulator of fimbrial expression in *K. pneumoniae* and its role on pathogenicity of this bacterium.

First, we showed that the Fur regulator binds to the Fur box identified on *kpfR* gene only in the presence of iron and that iron deprivation downregulates the expression of *kpfR*. Thus, we propose that the Fur regulator acts as a transcriptional activator of the *kpfR* gene. Since *kpfR* encodes a regulator of fimbriae transcription, Fur exerts indirect regulation of type 1-like fimbriae by positively regulating *kpfR*. An indirect role of Fur in expression regulation of fimbriae is not uncommon and has been described before. Wu et al. showed that the Fur indirectly controls the expression of type 3 fimbriae by positively regulating *mrkHI*, which are involved in the regulation of type 3 fimbriae *mrk* gene cluster (21).

The *K. pneumoniae* strain mutant for *kpfR* gene exhibited phenotypic changes, among which macroscopic changes on colony morphology, a slight dispersion on blood agar plates, and increased production of fimbrial structures. In fact, the mutant strain presented up-regulation of the type 1-like fimbriae gene clusters *fim* and *kpf*. Because of this increased production of type 1-like fimbriae, the mutant strain exhibits a greater ability to agglutinate yeast cells and to form biofilm. Taken together, these results point KpfR as a repressor of fimbriae transcription. Interestingly, this hyperfimbriated phenotype seems to provide the mutant strain a kind of rudimentary motility. Although *K. pneumoniae* is recognized as immobile, the finding of a rudimentary motility in this bacterium is not an unprecedented result. Carabarin-Lima et al. reported the presence of flagellar genes on a *K. pneumoniae* strain isolated from nosocomial infections and described a swim-like motility phenotype mediated by flagella in these clinical isolates (28). Besides, the *K. pneumoniae* genome contains the *flk* gene that encodes a poorly characterized regulator of flagella biosynthesis. It remains to be clarified whether the rudimentary movement observed in the mutant strain is due to the excess of type 1 fimbriae production, or if the KpfR regulator plays some direct role in regulating the expression of flagellar genes.

One of the phenotypic changes observed in the *kpfR* mutant strain was a drastic reduction in the production of capsular polysaccharide as opposed to the higher expression of fimbriae structures. Reports in the literature describe an inverse effect between the production of fimbriae and capsules, and also suggest that capsule production can inhibit the expression of fimbriae and interferes in fimbriae functionality (2, 29-31). In this respect, it is suggested that a possible coordinated regulation between fimbriae and capsules occurs through an environmental stimulus not yet known. For instance, Schwan et al. showed that adherence of uropathogenic *E. coli* to mannose receptors on bladder epithelial cells via type 1 fimbriae triggers a cross-talk that leads to the down-regulation of capsule production (32). Regarding *K. pneumoniae*, a possible cross-regulation between the capsule and fimbrial expression has been speculated, although not yet proven (9, 31, 33). A clue to this apparent cross-regulation of capsule and fimbriae expression in *K. pneumoniae* came from the work of Huang and coauthors (34). These authors showed that the deletion of the *cps* gene cluster, responsible for capsular polysaccharide biosynthesis, inhibits the expression of genes encoding type 1 and type 3 fimbriae. Corroborating these reports, our findings indicate KpfR as a repressor of fimbriae expression which, when knocked out, leads to activation of fimbriae expression and inhibition of the production of capsules in *K. pneumoniae*.

Although the hyperfimbriated phenotype yielded advantageous in *in vitro* coculture assays – as revealed by the greater efficiency of the mutant strain in adhering, invading, and replicating within eukaryotic host cells –, the increased expression of fimbrial structures proved to be detrimental during *in vivo* colonization in the murine model of urinary tract infection. In fact, our *in vivo* experiments showed that the *kpfR* mutant strain was unable to form intracellular bacterial communities (IBCs), exhibited lower titers in the bladder, and was quickly eliminated by the host. Therefore, by negatively controlling the expression of fimbriae and preventing the bacteria from having a hyperfimbriated phenotype, the KpfR regulator protects *K. pneumoniae* from being recognized and eliminated by the host immune system.

Our results with KpfR repressor bear similarities, but also striking differences, with FimK, another transcriptional regulator of fimbriae well characterized in *K. pneumoniae* (7, 19, 35). Unlike FimK, which has an EAL domain in its carboxy-terminal region (7, 19), KpfR does not have such a domain and, therefore, its mechanism of action should not involve the second messenger cyclic-di-GMP. The effects of KpfR and FimK on repressing the expression of type 1-like fimbriae emphasize the role of both regulators as inhibitory factors of biofilm formation. Also, both *kpfR* and *fimK* knockouts result in hyperfimbriated mutant bacteria (7) and reduced capsule production (35). However, contrary to that observed in *fimK* mutant strain, the *kpfR* mutant strain did not have its virulence increased in murine UTI model. FimK attenuates virulence on the urinary tract since lack of *fimK* resulted in higher bacterial titers, increased numbers of IBCs and enhanced virulence in the murine urinary tract (7). In contrast, KpfR seems to be essential for virulence in the urinary tract, because the loss of an active KpfR resulted in lower bladder titers, inability to form IBCs and attenuated effect on urinary tract virulence. A probable explanation for this is that in the absence of KpfR there is an induction of expression of two gene clusters of type 1-like fimbriae (*kpf* cluster and *fim* cluster), whereas FimK regulates only the *fim* gene cluster. Interestingly, the detrimental effect of FimK is restricted to the urinary tract, since in the respiratory tract it has a potentiating effect on virulence (35). Thus, both FimK and KpfR are repressors of fimbriae expression in *Klebsiella pneumoniae*. However, while FimK seems crucial for virulence in the respiratory tract, the KpfR revealed to be critical for UTI.

The loss of resistance and ability of the *kpfR* mutant strain to survive in the host can be attributed to its hyperfimbriated phenotype, which has made it to be readily recognized by the host defenses and becomes more efficiently and rapidly eliminated *in vivo*. In fact, while adherence is a crucial step for the pathogen to establish contact with the mucosa and invade and colonize the host tissue, pathogen-host interaction activates host antimicrobial defense, and a key event in this process is the bladder epithelium stimulation following bacterial adherence, which results in cytokines production by the epithelial cells through activation of host Toll-like receptor 4 (36). As stated by Song & Abraham, several Toll-like receptors have been identified on epithelial cells of the bladder which mediate a powerful immune response (37). Studies have shown that *K. pneumoniae* type 3 fimbriae can stimulate an oxidative response in neutrophils (25) and that the expression of fimbriae by *K. pneumoniae* increases their binding to phagocytes and triggers the phagocytosis, thus leading to bacterial killing (24).

In summary, pathogens employ survival strategies when in hostile environments, and to accomplish that they control the expression of virulence factors as needed. The expression of fimbriae represents an important factor of bacterial pathogenicity and, in this sense, it is of fundamental importance to understand the mechanisms that regulate the expression of this virulence factor. Herein, we described the role KpfR, a new transcriptional repressor of fimbrial expression hitherto not characterized, in the *K. pneumoniae* pathogenicity. By negatively controlling the expression of fimbriae, KpfR prevents *K. pneumoniae* from having a hyperfimbriated phenotype and from being eliminated in a murine model of urinary tract infection. Further investigations of the *kpfR* mutant strain on a murine model of respiratory tract infection will help to further elucidate whether loss of KpfR has an opposite effect according to distinct host niches.

## MATERIALS AND METHODS

### Bacterial strains, eukaryotic cell lines and growth conditions

For this study, we used *Klebsiella pneumoniae* strain #8 (UKP8), a clinical strain isolated from a human patient with urinary tract infection. Bacterial cells were routinely grown in Lysogeny Broth (LB; Bertani, 2004) at 37°C with shaking at 200 rpm, or on LB agar plates in static cultures. Bacterial growth was monitored by measuring the optical density of the cultures at a wavelength of 600 nm (O.D._600nm_), using the GeneQuant Spectrophotometer (GE Healthcare). For growth under iron-replete and iron-limiting conditions the LB medium was supplemented with FeSO_4_ (Sigma-Aldrich) or the iron chelator Dipyridyl (Sigma-Aldrich), respectively.

For coculture assays, we used the human bladder epithelial cell line T24, purchased from the Bank of Cells of Rio de Janeiro (BCRJ, cell line with code number 0231). T24 cells were cultivated at 37°C in an atmosphere of 95% relative humidity at 5% CO_2_, in McCoy’s 5A medium (ThermoFisher®) with 10% fetal bovine serum (FBS) (ThermoFisher®) and subcultivated twice a week at a ratio of 1:8.

### Identification of regulatory binding sites on *kpfR* gene promoter

Bioinformatics analyses were performed to identify putative regulatory binding sites on *kpfR* promoter. These analyses were conducted on the genomic sequence of *kpfR* on *Klebsiella pneumoniae* strain ATCC 700721/MGH 78578 (GenBank accession number CP000647.1). To identify Fur-binding sites on *kpfR* gene we employed a theoretical approach (38) previously adapted to *K. pneumoniae* (39). To identify the transcription start site and Sigma factor positions on the promoter region of *kpfR* gene we used the web-based programs *Neural Network Promoter Prediction* (available at http://www.fruitfly.org/seq_tools/promoter.html) and *BPROM-Prediction of Bacterial Promoters* (available at http://www.softberry.com/berry.phtml?topic=bprom&group=programs&subgroup=gfindb, (40), respectively. To determine whether *kpfR* cotranscribe with the genes from *kpf* cluster, reverse-transcription-PCRs were performed using cDNAs synthesized from total RNA extracted from *K. pneumoniae* and primer pairs spanning *kpfR* to *kpfA* and *kpfA* to *kpfD* genes (see Table S1 in the supplemental material). The resulting amplicons were analyzed by agarose gel electrophoresis. Primers were designed using *Primer3 version 0*.*4*.*0* web-program (available at http://bioinfo.ut.ee/primer3-0.4.0/, (41).

Complementary oligonucleotides containing the sequences of the putative Fur boxes were annealed to form double-stranded DNA probes. These DNA probes were cloned into high copy number pGEM®-T Easy vector (Promega) and the recombinant vectors were used in Fur titration assay (FURTA) and DNA Electrophoretic Mobility Shift Assay (EMSA), in order to validate the Fur protein interactions with the putative Fur boxes.

FURTA was performed with *Escherichia coli* strain H1717, kindly provided by Professor Dr. Klaus Hantke (University of Tübingen, Germany). *E. coli* H1717 carries the *lacZ* reporter gene under control of the Fur-regulated *fhuF* gene promoter. When transformed with multicopy plasmids cloned with functional Fur box, H1717 strain will appear red on MacConkey lactose agar plates (Lac^+^ phenotype) because the high number of newly introduced Fur boxes will titrate the Fur repressor from *fhuF*::*lacZ* fusion, thus releasing the transcription of *lacZ*. On the other hand, if H1717 strain is transformed with a vector cloned with a non-functional Fur box, the Fur repressor will remain bound to the *fhuF* promoter region and the the *lacZ* reporter gene is not expressed, rendering colorless *E. coli* H1717 colonies on MacConkey lactose agar plates (Lac^−^ phenotype).

FURTA was conducted as previously described (39). *E. coli* strain H1717 was transformed with pGEM®-T Easy vector cloned with the DNA probe containing the sequences of the putative Fur box identified on *kpfR* gene. The transformants were then plated onto MacConkey lactose agar containing 100 μg/ml ampicillin and 100 mM of iron sulfate (FeSO4) (Sigma). After 18 h of incubation at 37°C, the functionality of the putative Fur box on *kpfR* was assessed according to the Lac +/– phenotype (i.e., the color of the H1717 colonies). *E. coli* strain H1717 transformed with circular pGEM®-T Easy vector alone (i.e., with no insert) was used as negative control, whereas H1717 strain transformed with vector cloned with the previously validated Fur box of the *K. pneumoniae entC* gene (42) was used as a positive control.

EMSA was conducted essentially as described by Gomes et al. (39). DNA probes for EMSA were obtained by PCR amplifying the pGEM®-T Easy vector cloned with the putative Fur identified on *kpfR* gene, using the universal M13 primers (39). The resulting PCR product of 285 base pairs was then used as probes on the binding reactions, along with the previously purified His-tagged recombinant Fur protein from *K. pneumoniae*. Reactions were performed with 500 ηM of His-Fur protein, previously equilibrated for 10 minutes on ice in 10 μl reaction volume containing 1x binding buffer (10 mM Tris, 50 mM KCl, 1 mM DTT, pH 7.5), 0.5 mM MgCl_2_, 0.5 mM MnSO_4_ and 2.5% (v/v) glycerol. Next, 50 ηg of the DNA probes were added and the mixture was incubated for 20 minutes on ice. EMSA was also performed under divalent cation-free conditions by adding EDTA to a final concentration of 2 mM in the above reaction mixture. Samples were loaded onto a 2% (w/v) agarose gel prepared with 1x Bis-Tris borate buffer containing 0.1 mM MnSO_4_. After 30 minutes of electrophoresis the gels were stained with ethidium bromide solution (0.5 µg/ml) for 15 minutes, and the DNA bands were visualized and recorded under a digital photodocumentation system (Biorad).

### Construction of the *kpfR* mutant strain

Knockout of the *kpfR* gene from *K. pneumoniae* UKP8 was constructed using the *TargeTron Gene Knockout System* (Sigma-Aldrich), according to the manufacturer’s instructions. This system is based on site-specific and not random disruption of the gene of interest by insertion of group II intron harboring a kanamycin resistance gene (*kan*^R^).

Firstly, a computer algorithm at TargeTron Design website (Sigma-Aldrich) was used to identify potential target sites for intron insertion on the *kpfR* coding region. The most efficient target site was selected, based on the lowest *E-value* predicted by TargeTron Design algorithm (see Table S2 in the supplemental material). The TargeTron Design algorithm also provided the primers used in PCR reactions to mutate (re-target) the RNA segment of the intron (see Table S3 in the supplemental material). The 350 pb PCR product was digested and ligated into chloramphenicol-resistant pACD4K-C vector provided by the manufacturer (Sigma-Aldrich). The resulting recombinant pACD4K-C vector was transformed into *Escherichia coli* DH5a by heat shock, and clones of the recombinant vector was obtained using *Wizard® Plus SV Minipreps DNA Purification kit* (Promega). The pACD4K-C vector contains a T7 promoter and, therefore, requires T7 RNA Polymerase to express the retargeted intron that will disrupt the target gene. Since *Klebsiella pneumonia* does not express T7 RNA Polymerase, it was necessary to use the ampicillin-resistant pAR1219 vector (Sigma-Aldrich) in co-transformations because this vector contains the T7 RNA Polymerase gene under the control of the IPTG-inducible *lac* UV5 promoter. Chemically-competent *K. pneumoniae* UKP8 cells were co-transformed with pAR1219 and the pACD4K-C recombinant vectors and cultured overnight in LB containing 100 µg/mL of ampicillin and 25 µg/ml of chloramphenicol. On the next day, culture was diluted 1:50 in fresh LB containing ampicillin and chloramphenicol, and cell growth was monitored until they reached O.D._600nm_ of 0.2. At this point, in order to induce intron expression and insertion, 1 mM (final concentration) of IPTG was added to the medium and the culture was incubated at 30 °C for 30 minutes. After induction, cells were harvested by centrifugation at 12,000 rpm for 1 minute, resuspended in antibiotic-free LB broth and incubated again for 1 h at 30 °C. The cells were then plated on LB agar supplemented with 25 µg/ml kanamycin and grown at 30 °C for 2-3 days. Knockout bacteria were selected from kanamycin-resistant colonies and the insertion of the intron RNA harboring *kan*^R^ gene was confirmed by PCR using primers flanking the target insertion site in the coding region of the *kpfR* gene (Table S1 in the supplemental material). Since the knockout is based on the insertion of *kan*^R^ gene on *kpfR*, the *K. pneumoniae* UKP8 mutant strain was renamed *kpfR*::*kan*^R^.

A complemented strain, named *kpfR*::*kan*^R^_comp_, was obtained by inserting the *kpfR* gene back into the mutant strain. For this, a DNA fragment comprising the entire coding region of *kpfR* plus approximately 550 pb of 3’ and 5’ flanking regions (Table S4 in the supplemental material) was inserted on pCR2.1-TOPO vector (Invitrogen) previously cloned with erythromycin-resistance gene. Chemically-competent *kpfR*::*kan*^R^ was transformed with the recombinant vector and platted on LB agar supplemented with 50 μg/ml erythromycin. Complemented strains was recovery by screening erythromycin-resistant colonies.

### Growth experiments and phenotypic assays

Initially, growth of wild-type UKP8 and mutant *kpfR*::*kan*^R^ strains was assessed to investigate whether the lack of KpfR compromises bacterial growth. Strains were separately inoculated into LB medium and grown until saturation (overnight) at 37 °C under shaking. The next day, culture was diluted 1:200 in fresh LB and the bacterial growth was monitored every 15 minutes by measuring the O.D._600nm_. Growth curves were constructed by plotting the O.D._600nm_ values against time.

Ten microliters of saturated cultures of UKP8, *kpfR*::*kan*^R^ and *kpfR*::*kan*^R^_comp_ were platted on blood agar (Trypticase Soy Agar plates supplemented with 5% sheep blood) during 18 h at 37°C to check for macroscopic differences in each strain’s morphology.

Motility assays were carried out in triplicate for each *K. pneumoniae* strains. In brief, 10 µL of bacterial cultures standardized at O.D._600nm_ of 0.4 was inoculated in the center of soft LB agar plates (prepared with 0,8% bacteriological agar) and incubated at 37°C for 120 h. After this, the extension of the colonies diameter was measured in centimeters.

Transmission Electron Microscopy (TEM) was carried out for visualization of fimbriae production by the *K. pneumoniae* strains. Bacterial cells were grown statically on LB agar plates at 37°C for 18 h and resuspended in 500 μl of PBS buffer. A droplet of the cell suspension was spotted on Formvar coated carbon-reinforced copper grid (400 mesh) and incubated at room temperature for 2 minutes. Grids were negatively stained using a drop of 1.25% phosphotungstic acid pH 6.5 (Sigma-Aldrich) filter sterilized through a 0.22 µm pore-size filter membranes (Millipore). The staining lasted 15 seconds and then the grids were air-dried on a piece of filter paper. Samples were examined and electron micrographs were obtained with a Zeiss LEO 906 Transmission Electron Microscope operated at 60 kV.

Biofilm formation assays were conducted in polyvinylchloride (PVC) microplates, following a protocol described elsewhere with minor modifications (43). This protocol is based on the ability of bacteria to form biofilm on PVC microplates and on the ability of crystal violet to stain bacterial cells but not PVC. Strains were grown until saturation at 37 °C under shaking. Bacterial cells from saturated cultures were harvested by centrifugation and resuspended in LB broth to a final concentration of 10^6^ cells/mL. 5 µL of this suspension were added on flat-bottom 96-well PVC microplates containing 150 μL of LB and incubated at 37°C for 1, 5 and 10 h under static conditions. After incubation, the culture medium was removed and the wells were gently washed twice with sterile water to remove unattached cells. Next, 25 μl of 1% crystal violet was applied in the wells and the plates were incubated for 15 min at room temperature. After incubation, the excess of dye was removed and the wells were washed with deionized water. Then, the remaining crystal violet staining the adherent cells was solubilized with absolute ethanol and the absorbance of the ethanol containing the eluted dye was measured at O.D._600nm_ in a spectrophotometer. The biofilm formation assays were repeated at least three times.

Yeast agglutination assays were performed to investigate the expression of fimbriae by the *K. pneumoniae* strains. Assays were conducted on Kline concavity slides, according to protocol previously described (31, 44). Bacterial strains grown on LB agar plates up to 72 h were resuspended in PBS buffer and the concentration was standardized at O.D._600nm_ of 0.6. Bacteria were then mixed with equal volume of 5% (wt/vol) suspension of *Saccharomyces cerevisiae* cells (Sigma-Aldrich) prepared in PBS buffer. The time required for agglutination and its intensity were documented. The agglutination of the yeast cells is specifically mediated by type 1 fimbriae since these fimbrae have great affinity for mannose, a highly abundant residues on yeast cell-surface. Therefore, the assays were also performed in the presence of 5% D-(+)-Mannose (Sigma-Aldrich) to confirm if the agglutination was indeed mediated by the type 1 fimbriae.

To visualize capsular polysaccharide production, UKP8, *kpfR*::*kan*^R^ and *kpfR*::*kan*^R^_comp_ strains were cultured under the same growth conditions, stained with India ink and analyzed by optical microscopy. The presence of capsule production is indicated by a negative staining area around the bacteria.

### Epithelial cell adhesion, invasion and intracellular replication assays

Coculture assays were performed as previously described (45). Saturated overnight cultures of UKP8 and *kpfR*::*kan*^R^ were inoculated on fresh LB medium and incubated at 37°C with shaking until they reached the mid-logarithmic growth phase (O.D._600nm_ of 0.4). Bacteria were added to an 80% confluent T24 cell monolayer seeded into 12-well plates at a multiplicity of infection (MOI) of 200 bacteria per host cell, and the infected monolayers were incubated at 37°C for three different time periods, specific for each assay. For adhesion assay, infected T24 cells were incubated for 30 minutes, washed three times with phosphate-buffered saline (PBS buffer) to remove non-adhered bacteria, and lysed by adding 0.1% Triton X-100 diluted in PBS. Serial dilutions of the lysate were plated on blood agar and incubated overnight at 37 °C for counting bacterial colony-forming units (CFUs). For invasion assay, infected T24 cells were initially incubated for 4 h to allow invasion of the bacterial strains into T24 cells. Plates were washed once with PBS buffer and incubated again for an additional 1 h in fresh culture medium containing the antibiotic gentamicin (25 µg/ml) to eliminate extracellular bacteria (both planktonic and adhered). The plasma membrane of host cells is impermeable to gentamicin. Therefore, after treatment with this antibiotic, only extracellular bacteria will be eliminated, while those present in the intracellular environment will be preserved. After treatment with gentamicin, the T24 cells were washed, lysed with Triton X-100 and serial dilutions of the lysate were plated on blood agar for counting of CFUs. For intracellular replication assay, infected T24 cells were initially incubated for 4 h to allow invasion of the bacterial strains. Plates were washed once with PBS buffer and incubated again for 24 h in fresh culture medium containing gentamicin at 10 µg/ml. Then, the T24 cells were washed, lysed with Triton X-100 and serial dilutions of the lysate were plated on blood agar for CFUs counting. The integrity of the T24 cells during the coculture assays was assessed by the standard exclusion protocol of trypan blue 0,4% (ThermoFisher®).

### Mouse model UTI

For mouse urinary tract infection analyzes were used 6-8 weeks-old female BALB/c mice (CEMIB, UNICAMP, Brazil). Infections were carried out by transurethral inoculation as previously described (46, 47). UKP8 and *kpfR*::*kan*^R^ strains were grown on LB agar plates at 37°C for 18 h and resuspended at a concentration of ∼5 x 10^8^ CFU/ml. Fifty microliters of the bacterial suspension were inoculated by transurethral catheterization in mice anesthetized intraperitoneally with ketamine and xylazine. Mice were euthanized at the indicated times, and urine and bladders were aseptically collected and processed for histology and CFU titration. Serial dilutions of urine and homogenized bladders were plated on blood agar and incubated overnight at 37 °C for counting of CFUs. Whole bladders were fixed in 10% neutral buffered formalin, embedded in paraffin, and 5 μm-thick sections were stained with hematoxylin and eosin (H&E).

Mice experimental procedures were conducted in accordance with guidelines of the Brazilian College for Animal Experimentation (COBEA) and received prior approval by the Ethics Committee on the Use of Animals in Research of Universidade São Francisco (Approval number 01.0226.2014).

### RNA extraction and RT-qPCR analysis

To investigate the expression of *kpfR* under iron-replete and iron-limiting conditions, *K. pneumoniae* strains were grown in LB broth at 37°C with shaking at 200 rpm until they reached the mid-logarithmic growth phase (O.D._600nm_ of 0.4). At this point, ferrous iron (100 μM of FeSO_4_, final concentration) or an iron chelator (100 μM of 2,2’-Dipyridyl, final concentration) were added and the cells were incubated for 1 h. After 1 h incubation, bacterial cells were harvested by centrifugation and the pellets were resuspended in *RNAprotect® Bacteria Reagent* for RNA stabilization. All culture conditions were performed at least twice. Control condition consisted of *K. pneumoniae* cells grown in LB medium without supplements.

The expression of fimbrial genes were also assessed on UKP8 and *kpfR*::*kan*^R^ strains grown on blood agar plate. For this, bacterial colonies from each strain were individually collected from the blood agar plates, and the cell pellets were resuspended in *RNAprotect® Bacteria Reagent* until the moment of the RNA extraction.

Total RNA from the abovementioned cell pellets was extracted using *RNeasy Protect Bacteria Mini Kit* (Qiagen), following the manufacturer’s protocol. An on-column DNase digestion with the RNase-free DNase Set (Qiagen) was performed to remove genomic DNA contamination in RNA samples.

To assess gene expression of the bacterial strains during coculture with T24 bladder epithelial cells, adhesion and invasion assays were conducted as previously described, the only exception that the T24 cells were seeded in 06-well plates. After the incubation periods, extracellular bacteria were removed by washing with PBS buffer and T24 cells were collected for the extraction of bacterial total RNA. RNA was extracted using Max™ Bacterial RNA Isolation kit (Ambion), following the manufacturer’s instructions. MICROBEnrich™ kit (Ambion) was applied, in order to promote the selective removal of total RNA from T24 cells, preserving bacterial RNA. Total RNA extracted was further treated with DNAse.

After treatment with DNAse, 0.5 to 1 μg of total bacterial RNA was used for cDNA synthesis by using the *ThermoScript™ RT-PCR System for First-Strand cDNA Synthesis Kit* (Invitrogen) according to the manufacturer’s instructions. The synthesized cDNAs were used in RT-qPCR reactions done in triplicates using the *Platinum® SYBR® Green qPCR SuperMix-UDG* Kit (Invitrogen) on *7300 Real-Time PCR System* equipment (Applied Biosystems). Data were normalized using *rho* and *recA* as endogenous genes, which encode transcription termination factor and recombinase A, respectively (39). The relative expression levels of the selected genes were calculated by the comparative critical threshold (ΔΔC_T_) method (48). GraphPad Prims 7.00 (GraphPad Software, Inc.) was used for the statistical analyses. Differences on the expression levels were evaluated by Student’s *t* test, and differences with *p*-values ≤ 0.05 were considered statistically significant. Primers used on RT-qPCR reactions are listed in Table S5 in the supplemental material and were designed using *Primer3 version 4*.*1*.*0* web-program.

## ACKNOWLEDGMENTS

This work was supported by the São Paulo Research Foundation (FAPESP), grants #2008/11365-1 and #2013/06042-7. AEIG and CSS were supported by scholarships from Coordination for the Improvement of Higher Education Personnel (CAPES, Ministry of Education of Brazil). TP was supported by a scholarship from São Paulo Research Foundation (FAPESP), grant#2013/13949-9.

## SUPPLEMENTAL MATERIAL

**Figure S1.**
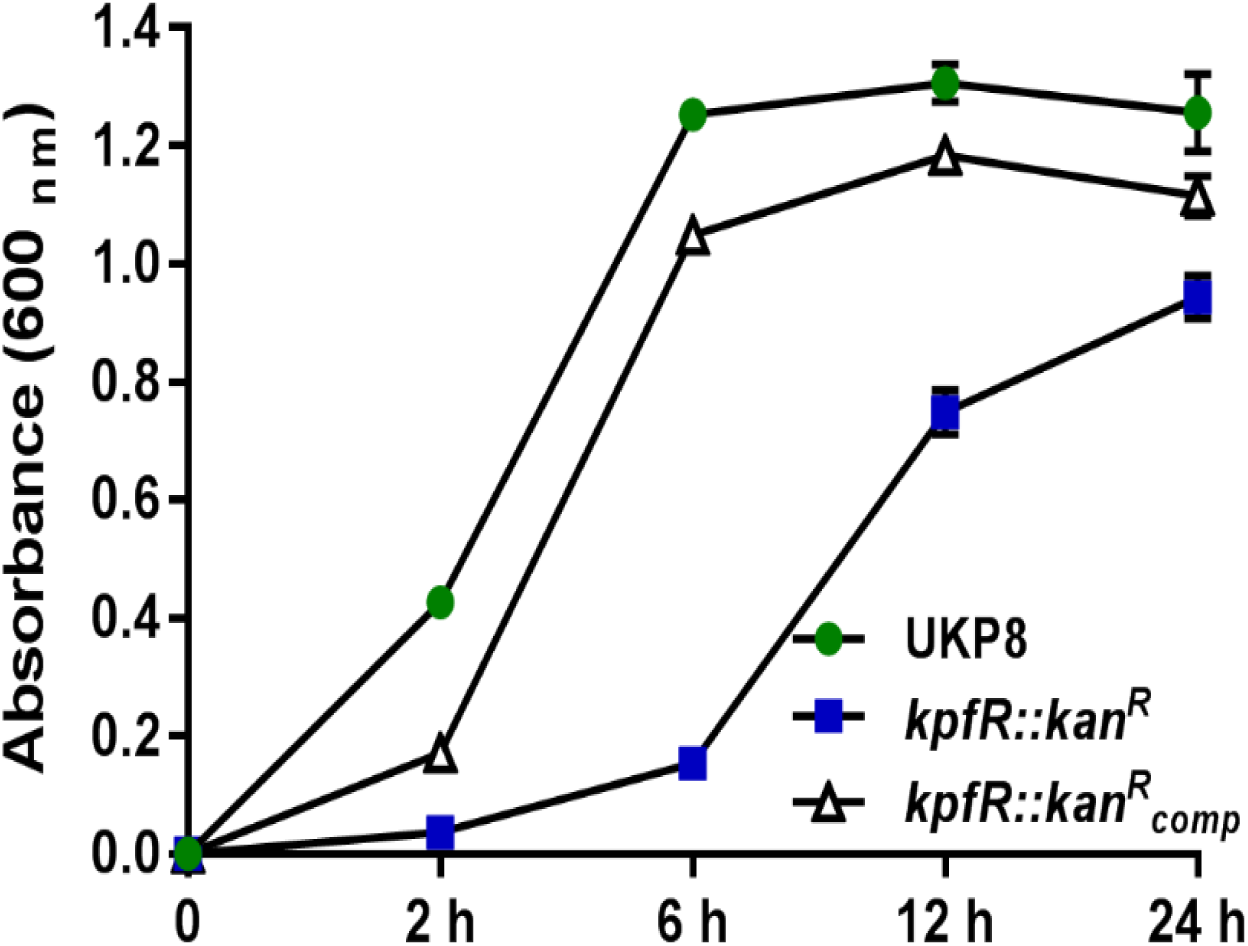
Growth curves of UKP8, *kpfR*::*kan*^R^ and *kpfR*::*kan*^R^_comp_ assessed by monitoring the optical density of the strains grown in LB medium at 37 °C with agitation. Despite the delayed entry into logarithmic (log) phase due to prolonged growth at lag phase, the mutant strain *kpfR*::*kan*^R^ exhibit a growth rate similar to that observed in the wild-type UKP8 on the log phase. The growth curve observed on the wild-type strain was reestablished on the complemented mutant strain *kpfR*::*kan*^R^_comp_.

**Table S1.**
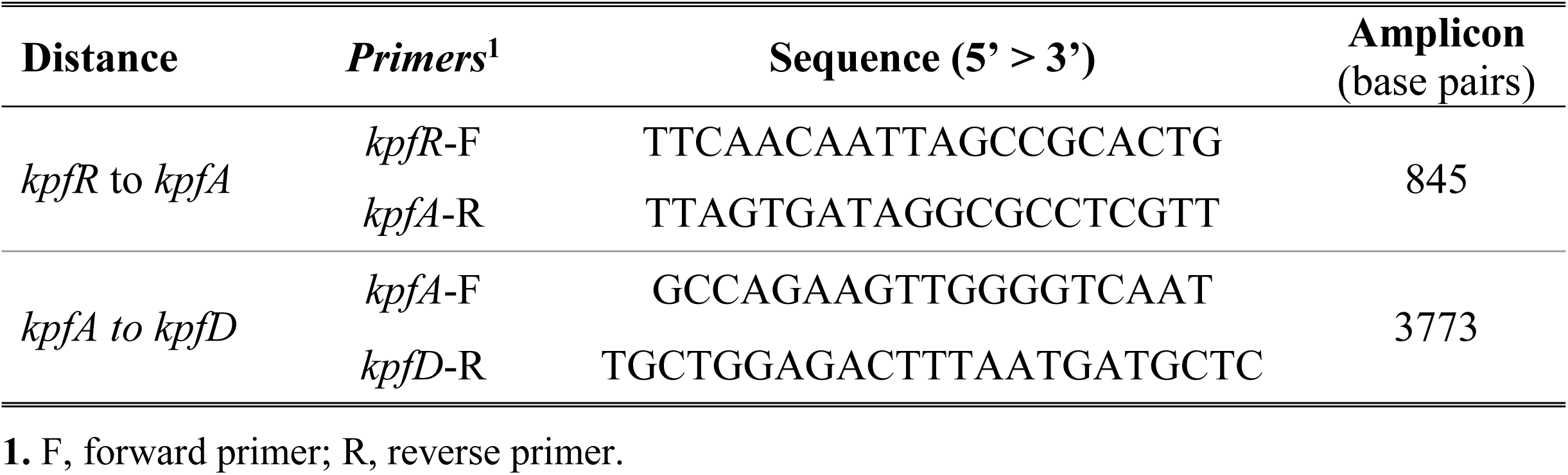
Primer pairs used in PCR reactions to confirm the polycistronic transcription of *kpfR* and *kpf* gene cluster. Primer pairs *kpfR*-F and *kpfA*-R were also used to confirm the insertion of the intron in the coding region of the *kpfR* gene.

**Table S2.**
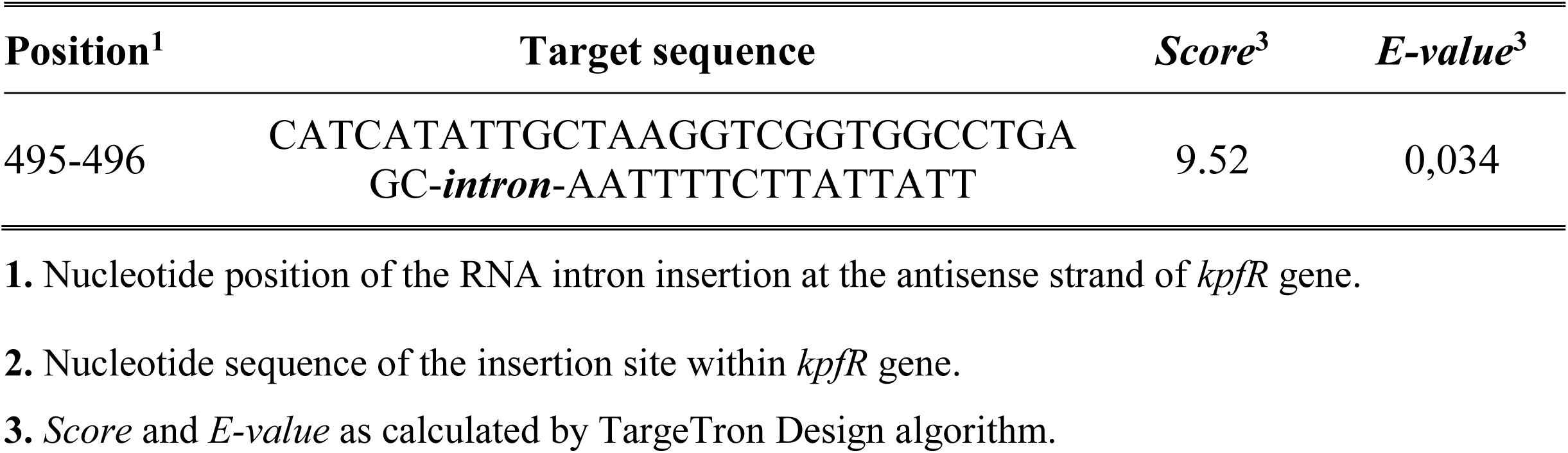
The most efficient insertion site on *kpfR* coding region predicted by TargeTron Design algorithm.

**Table S3.**
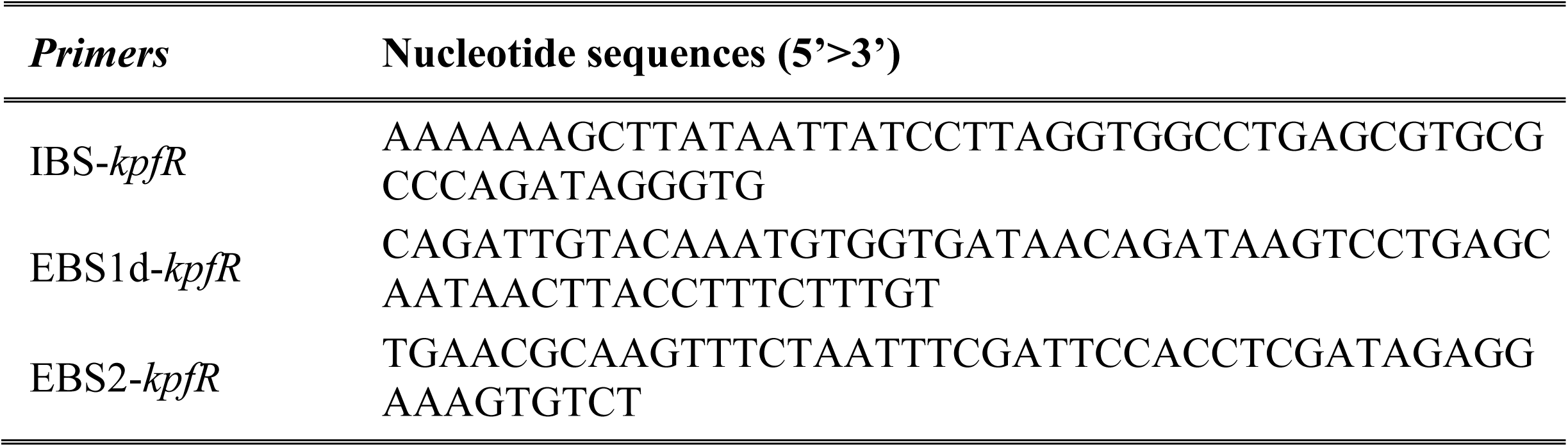
Nucleotide sequence of the primers IBS-*kpfR*, EBS1d-*kpfR* and EBS2-*kpfR* primers generated by TargeTron design algorith. These primers were used in PCR reactions to retarget the RNA segment of the intron.

**Table S4.**
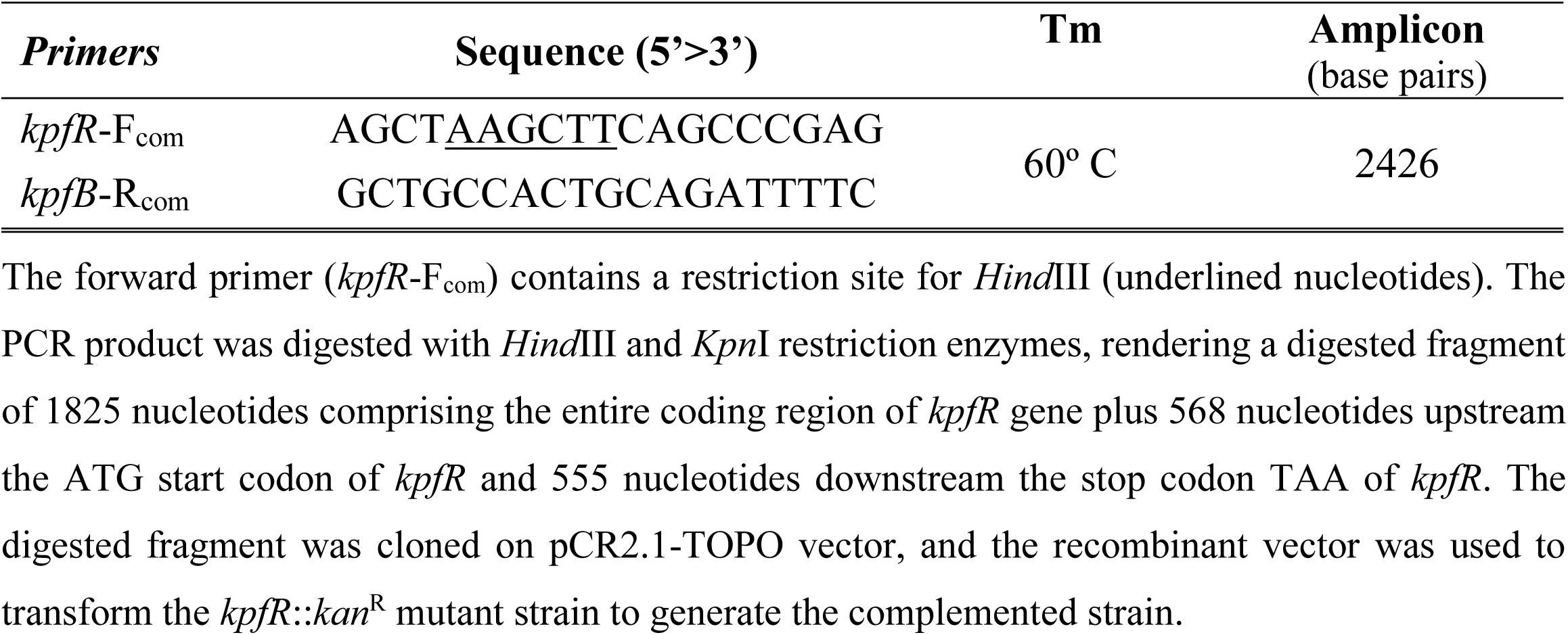
Primer pairs used to amplify the entire *kpfR* coding region plus 3’ and 5’ flanking regions to generate the complemented strain.

**Table S5.**
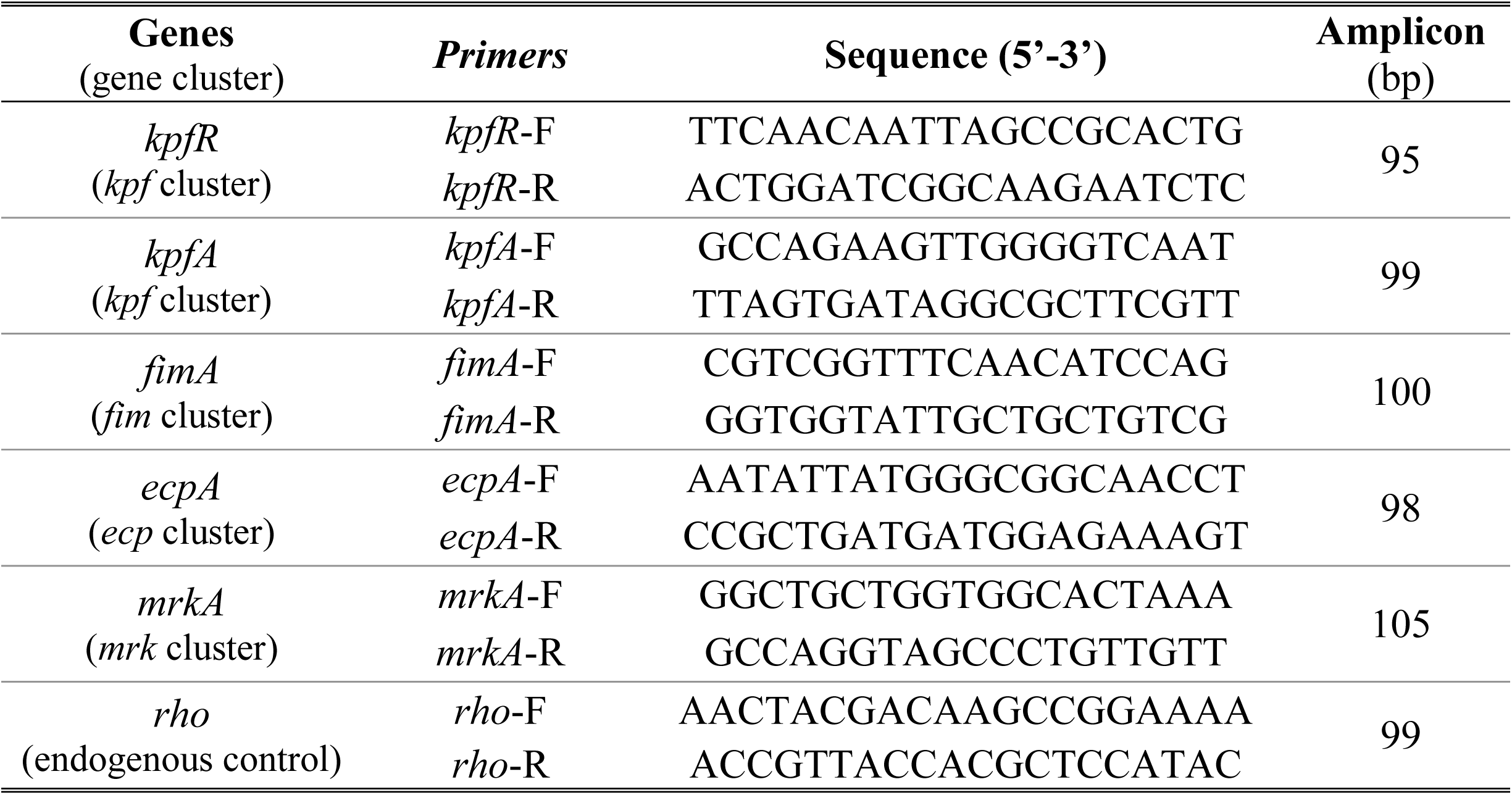
Primer pairs used on RT-qPCR reactions. Primers were designed using *Primer3 version 4*.*1*.*0* web-program, in order to present 60 °C of annealing temperature and amplicon sizes ranging from 95 to 105 base pairs (bp).

